# The pathway to resolve dimeric forms distinguishes plasmids from megaplasmids in Enterobacteriaceae

**DOI:** 10.1101/2024.04.05.588136

**Authors:** Florian Fournes, Manuel Campos, Jean Cury, Caroline Schiavon, Carine Pagès, Marie Touchon, Eduardo PC Rocha, Philippe Rousseau, François Cornet

## Abstract

Bacterial genomes contain a plethora of secondary replicons of divergent size. Circular replicons must carry a system for resolving dimeric forms, resulting from recombination between sister copies. These systems use site-specific recombinases. Among these, the XerCD recombinase resolves dimers of chromosomes and certain plasmids using different controls. We have analyzed the dimer resolution functions in enterobacterial secondary replicons and show that, in addition to the main chromosomes, XerCD is preferentially used by small plasmids and by the largest secondary replicons, megaplasmids and secondary chromosomes. Indeed, all replicons longer than 250 kb host an active XerCD recombination site. These sites, in contrast to those of small plasmids, use the same control as chromosomes, coupled to cell division by the FtsK protein. We conclude that a chromosome-like mode of dimer resolution is mandatory for the faithful inheritance of large plasmids and chromids, its acquisition being a prerequisite for the genesis of secondary chromosomes from plasmids.

## Introduction

Beside a unique main chromosome, bacterial genomes contain a plethora of secondary replicons differing in fate, composition and size (1–8). These are found in most bacterial genomes and carry adaptive traits along with functions for their own maintenance and dissemination. Most of them are mobile genetic elements that can transfer between strains and species, bringing for example the acquisition of symbiosis or pathogenic power and resistance to antibacterial compounds (9–11).

Secondary replicons differ in size, copy number and the maintenance and transfer functions they encode, allowing the definition of replicon categories (1, 12). Small ‘high-copy-number’ plasmids are less than 25 kilobases (kb) long. Moderate to low-copy-number (LCN) plasmids are larger, most of the time about 100 kb in enterobacteria, and frequently encode the complex machinery for their self-transfer by conjugation. The term megaplasmids was proposed for notably large LCN plasmids that show no important chromosome-like feature (5, 7). They frequently have mosaic structures, carrying evidence of plasmid co-integration and of extensive gene transfer (13). Lastly, chromids and secondary chromosomes, often referring to the same replicon categories, are large replicons, typically above 500 kb, found in the different strains of a specie, thus becoming part of its core genome (6, 7, 14). They carry core genes and appear more adapted to their host than plasmids and megaplasmids since they display nucleotide composition and codon usage closer from the main chromosome.

Circular replicons should contain at least one system to resolve dimeric forms (12, 15, 16). Dimers or higher multimers arise from homologous recombination between sister copies and lower the number of plasmid copies, hindering stable inheritance. They are resolved by site-specific recombination.

Resolution systems have been mainly studied in some model HCN plasmids and in chromosomes. Chromosomes use the XerCD recombinase, activated by the FtsK DNA translocase, to recombine the XerCD recombination site (*xrs*) they carry, called the *dif* site (fig 1A). *Xrs* are short pseudo palindromic sequences (28 to 30 bp) imperfectly conserved, consisting of two 11 bp binding sites for XerC and XerD, respectively, separated by a 6 to 8 bp central region (fig 1B). The terminal region of the chromosome (*ter*), where the *dif* site lies, carries numerous *matS* sites recognized by the MatP protein (fig 1A and B) (17). MatP delays *ter* segregation and tethers sister *ter* to the division septum to which the FtsK DNA translocase is linked. FtsK segregates sister *ter* following the orientation dictated by the KOPS motifs, ending at *dif*, then activates XerCD recombination in the case sister chromosomes are dimeric (fig 1; fig S1A; (12, 18)). Some model HCN plasmids, i.e. ColE1 and pSC101, also contain *xrs* and use XerCD. In this case, recombination does not depend on FtsK but requires 200 bp-long specific sequences (called accessory sequences, AS) adjacent to the XerC-binding site of the *xrs* (fig 1B). AS are recognized by specific proteins, either the ArgR (plasmid ColE1 and relatives) or ArcA (plasmid pSC101) and the PepA protein in both cases. These proteins form a nucleoprotein complex containing pairs of ASs only when present on the same molecule, ensuring recombination resolves dimers better than creating them (fig S1B; (16)). ArgR and ArcA bind cognate sites at defined position in ASs whereas PepA does not bind recognizable DNA motif but is attracted by either ArgR or ArcA (19).

**Figure 1:**
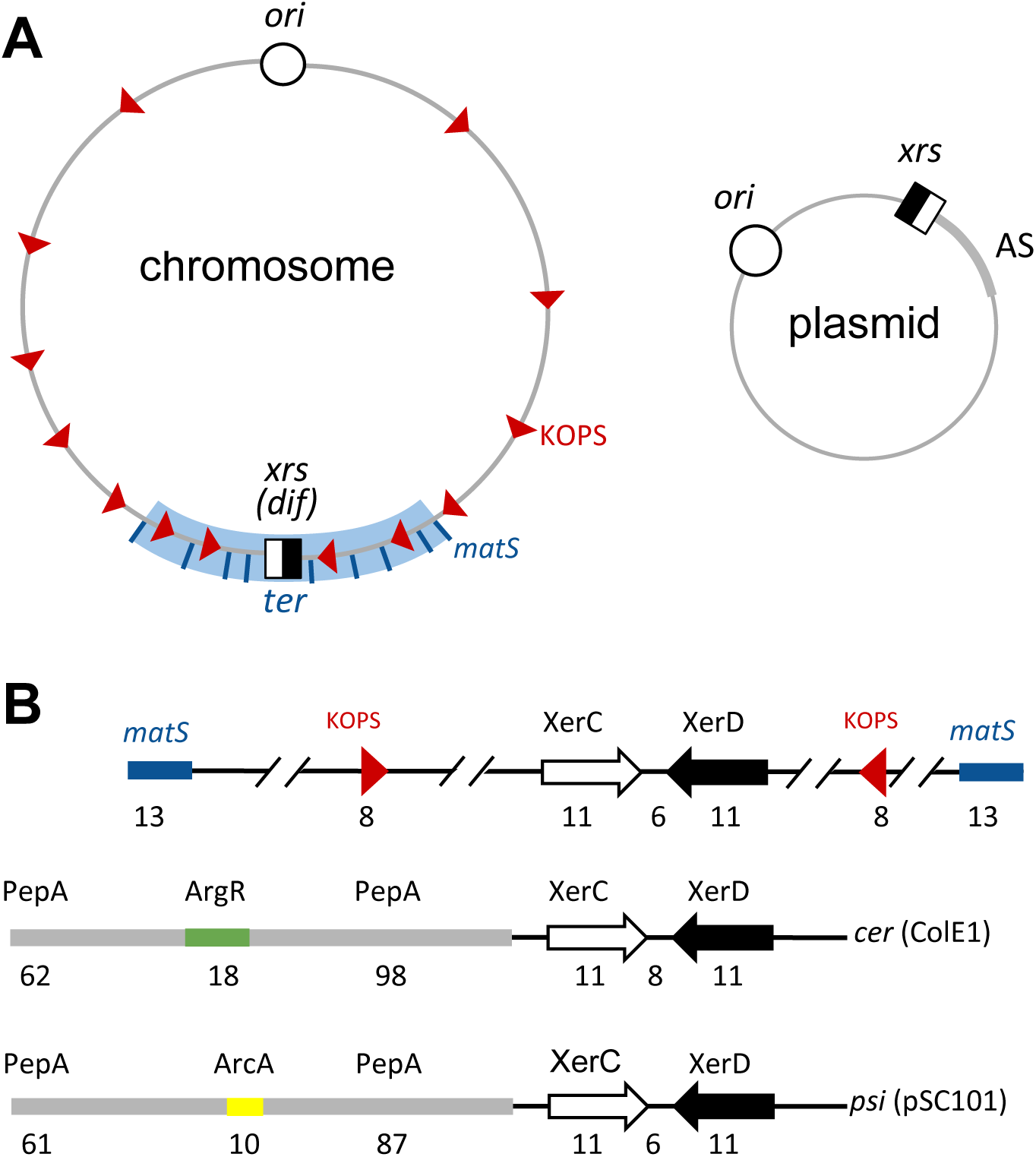
Xer recombination sites. A) Replicon maps showing *xrs* (black and white squares) with relevant DNA elements involved in the control of recombination: KOPS (red arrowheads), *matS* (blue lines) in the case of chromosomes (left) and accessory regions (AS, grey) in the case of small multicopy plasmids (right). B) Local organization around *xrs* in the case of the chromosome (*dif* site, top line) and the two model plasmids ColE1 (*cer* site, middle) and pSC101 (*psi* site, bottom line). Binding sites for XerC and XerD (open and black arrows, respectively) ArgR (orange) and ArcA (yellow) are shown. PepA binds with poor specificity in the regions surrounding the ArgR and ArcA binding sites. Sizes of the different elements are indicated in bp.

Here, we asked how the two described controls of dimer resolution are distributed in secondary replicons. We analyzed enterobacterial secondary replicons for their dimer resolution mode. We show that replicons encode site-specific recombinase and/or possess *xrs* depending on their size. HCN plasmids tend to harbor *xrs* associated with AS whereas LCN plasmids tend to encode their own recombinases. Strikingly, the larger replicons show a tendency to possess *xrs* which increase with their size and become a rule above 250 kb. We analyzed the R27 plasmid and show that it possesses an FtsK- activated *xrs* able to resolve its dimeric forms. Chosen *xrs* cloned from large replicons also recombined in an FtsK-dependent manner. We further show that KOPS biases and *matS* sites arise on the largest replicons. We conclude that acquiring FtsK-controlled dimer resolution is compulsory for replicons to grow significantly over the size of LCN plasmids.

## MATERIAL AND METHODS

### Strains, plasmids and standard procedures

DNA and *E. coli* strains manipulations and constructs all followed standard procedures. Plasmid R27 was a gift from Carlos Balsalobre (U. Barcelona, Spain). From the R27 sequence (20), gene R0101 was identified with the help of Laurence Van Melderen’s laboratory (U. Bruxelles, Belgium) as encoding a potential toxin. It was deleted with the neighbor gene, R0102, and replaced by the *lacI* gene, yielding our ’wt’ R27. Gene R0183 encodes a potential Y-recombinase and was replaced by an FRT-Kn-FRT cassette subsequently resolved to a single FRT site using the FLP recombinase. A potential *xrs* was identified as the 5’- AGTACATATACCAAAGATTATGTTAAAT sequence. This region was sequenced, allowing to correct the C at position 11 to an A. The corrected *xrs* was subsequently cloned in pUC57 backbone and used. Plasmids for recombination monitoring were constructed using PCR assembly to first obtain *xrs*-Kn module, which was then inserted in pUC57-derivatives already containing *xrs*, resulting in *xrs*-Kn-*xrs* cassettes.

Strains used for recombination assays are derivatives of DS941 (21) rendered Δ(*xerC*)::Cm, Δ(*pepA*)::Kn or Δ(*ftsK*C)::Tc. Strains were transformed by plasmids carrying directly repeated *xrs*, grown overnight on antibiotic-containing plates. Ten colonies were pick-up and grown before plasmid extraction and analysis on 1% TAE agar gels.

R27 loss was measured in derivatives of strain LN2666 (22) first rendered Δ(*lacI*)::Sp, then either Δ(*xerC*)::Cm, Δ(*xerD*)::Kn or *ftsK^ATP^*^-^-Cm by standard procedures. Strains containing R27 derivatives carrying the *lacI* gene were cultivated overnight in L broth plus tetracycline, then diluted twice per day and grown in serial cultures without tetracycline for three days, corresponding to 60 generations, before plating on X-Gal-containing plates. The ratio of blue colonies (R27 loss) on total is reported with standard deviation of three independent experiments.

### Protein purification and *in vitro* reactions

Proteins purifications were performed as described in (23) for *E. coli* XerC and XerD recombinases and in (24) for the tαβγFtsKC protein.

For electrophoretic mobility shift assays, radiolabeled 28-bp DNA fragments containing *xrs* were incubated with increasing concentration of XerC and/or XerD proteins (from 0.2 mM to 0.8 mM), as reported in and visualized using Typhoon-Trio-GE.

*In vitro* recombination experiments were mainly performed as previously reported in (25). Briefly, 300ng of plasmid DNA containing *xrs*-Kn-*xrs* cassette were incubated with final concentrations of 0.3mM of XerC and XerD proteins and 0.5mM of tαβγFtsKC, in a buffer containing 25 mM TrisHCl pH7.5, 10mM MgCl2, 0,1 % PEG8000 and 2.5mM ATP, and incubated for 2h at 30°C. Reactions were stopped using a buffer containing 10% SDS and 2mg/ml of proteinase K and then analyzed by 0.8% agarose gel electrophoresis.

### *xrs* sites and AS detection

Site-specific recombinases were searched using HMM profiles. To search for *xrs*, we started from the few *xrs* functional characterized (sup file 2), searched by homology using BLASTN then checked manually the retrieved sequences applying some rules : (i) the internal parts (first 5 bp) of the XerC and XerD binding sites are the most conserved and most often form a palindrome; (ii) the XerD binding site is better conserved than the XerC-binding site; (iii) the central region is 6 to 8 bp long and its sequence is not conserved. The retrieved *xrs* were added to the search until no additional *xrs* was retrieved. A total of 518 *xrs* were found, of which 183 unique sequences (sup file 1).

As the previously characterized ASs are poorly conserved, we relied on searching potential ArcA or ArgR binding sites at the same positions as in known AS. Retrieved AS were then used for homology search and this procedure was repeated until no new AS was retrieved. Most AS contained an ArgR binding site, only four harboring one for PepA. The most divergent examples are shown in sup. file 2.

### KOPS detection

We identified KOPS with the regular expression function from the built-in Python package re, matching the pattern “GGG[ACGT]AGGG” and its reverse complement ‘CCCT[ACGT]CCC”.

For representation analysis, we used R’MES (27) to calculate a score for the motif GGGNAGGG with a maximal markov chain length (l=6) and considered, as recommended, the values of -3 and 3 as threshold for under- and over-represented cases, respectively.

For skew analysis, KOPS positions were translated into angles after a circular shift of the plasmid DNA sequence so as to place the *xrs* site at the middle of the sequence. The angle of the *i*^th^ instance of the motif, *θ*_*i*_, is calculated as:

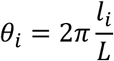

where *l*_*i*_ is the position of the *i*^th^ instance of the motif along the re-oriented sequence and L the length of the plasmid sequence.

KOPS clustering was evaluated with the resultant length of the set of angles formed by all instances along the plasmid DNA sequence. The square of the resultant length is defined as:

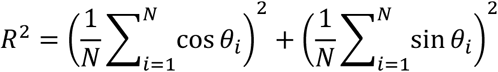

where *θ*_*i*_ is the angle of the *i*^th^ instance and N the number of instances

The statistical significance of clustering was estimated using the Rayleigh test included in the circstats module of the Astropy Python package (26). Low KOPS occurrences limit our ability to detect clustering in small plasmids. We estimated the statistical power to fall below 0.5 for plasmids with less than 15 KOPS occurrences.

### *matS* sites detection

The 23 *matS* of the chromosome of *E. coli* K12 MG1655 were used to construct a position-specific score matrix (PSSM). The threshold score to define a site as a bona fide *matS* was defines so as to selectively detect all 23 *matS* sites previously identified on *E. coli* K12 MG1655 chromosome (17).

To analyze *matS* representation, 1000 random sequences were generated for each plasmid using a custom Python script (rnseq.py) that extend the DNA sequence by matching their 4-mer distribution of each plasmid. The number of *matS* sites in these random sequences thus represents the expected number of *matS* sites under a Markov model with a Markov chain of length 4. The distribution of *matS* counts in these recoded sequences was effectively approached by a Poisson distribution used to estimate how (un)likely it is to observe the actual *matS* counts in real plasmid sequences as:

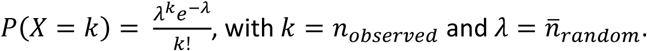

## Results

### *xrs* are present on plasmids depending on their size

To predict how secondary replicons cope with dimer resolution, we performed a survey of the site-specific recombinases they encode and a search for XerCD recombination sites (*xrs*) they carry (Material and Methods). We used a collection of 981 Enterobacteriaceae secondary replicons previously annotated for other maintenance functions (28). These replicons ranged from 1.3 to 794 kb in size and showed a bimodal size distribution: 25% were less than 25 kb and harbored replication functions suggesting either rolling circle-type replication or theta-type replication with high copy number per cell, while the others were centered around the 50 to 150 kb range (half of the plasmids) and predicted to be low copy-number from their replication functions (fig 2 lane A). Only 10% of the replicons were larger than 200 kb and 7 larger than 500 kb.

**Figure 2:**
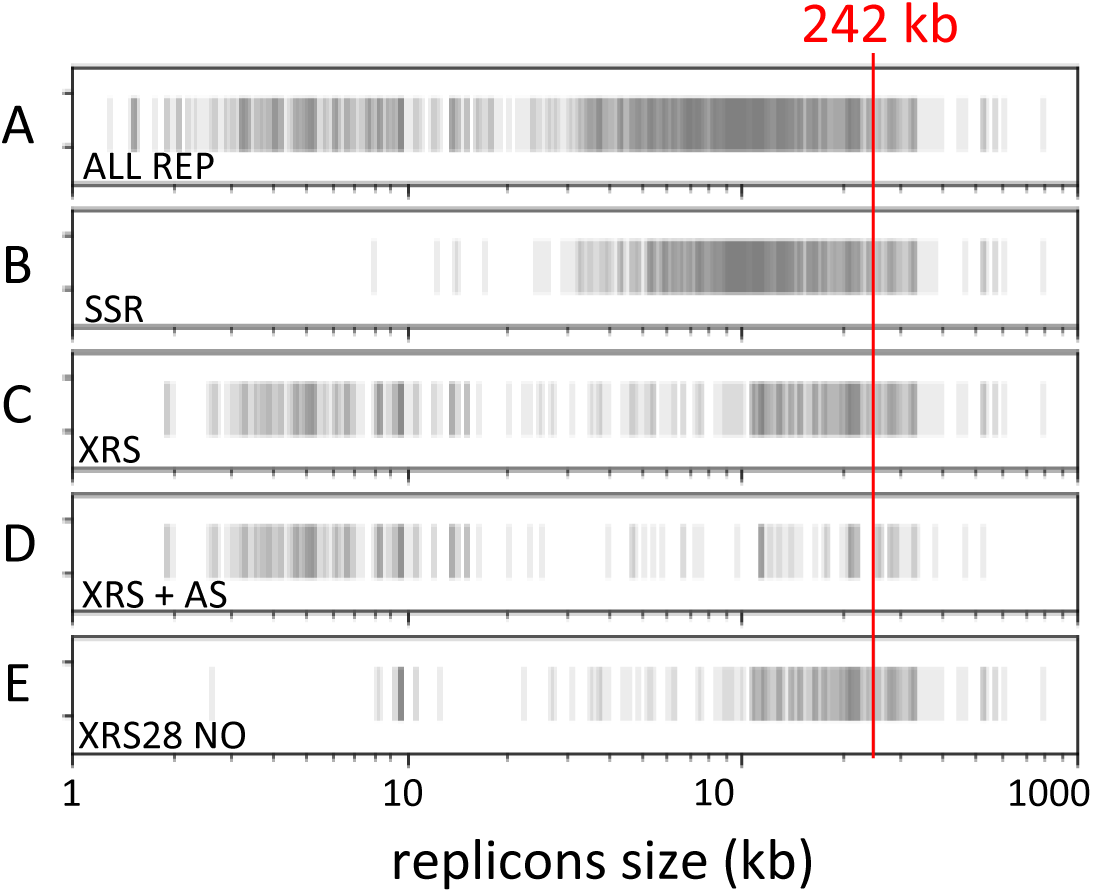
Dimer resolution systems depend on replicon size. Eventplots depending on replicon size (x-axis). A: all 981 replicons; B: presence of at least one site-specific recombinase; C: presence of at least one potential *xrs*; D: presence of at least one potential *xrs* flanked by a potential AS next to the XerC binding site; E: presence of at least one potential 28 bp long *xrs* (6 bp long central region) devoid of recognizable AS. The size above which all replicons harbor at least one 28 bp-long xrs devoid of AS is indicated in red.

Site-specific recombinases were frequent in plasmids larger than 25 kb and tended to be absent in smaller plasmids (fig 2 lane B). Small plasmids tended to harbor *xrs* (70% of the plasmids smaller than 25 kb; fig 2 lane C) while medium-sized plasmids (30 to 100 kb) tended to have no *xrs*. Strikingly, most plasmids larger than 115 kb and all replicons above 248 kb had at least one *xrs*.

*Xrs* may recombine depending on accessory sequences (AS) and proteins whatever their size : 28, 29 or 30 bp. In contrast, only 28 bp *xrs* were shown to recombine depending on FtsK (16, 29, 30). Of the eight AS previously reported, only two pairs displayed weak yet significant homology (Material and Methods). We thus relied on the presence of ArgR or ArcA binding sites at the correct distance from the *xrs* (fig 1B) in addition to homology search of ASs (Material and Methods). All *xrs* present on small plasmids were associated with an AS whereas *xrs* present on large replicon tended to be devoid of AS (fig 2 lane D). In contrast, 28 bp *xrs* not associated with an AS were almost restricted to large replicons (fig 2 lane E), suggesting that these replicons use FtsK-dependent recombination to resolve dimers. Taken together, our analysis suggests three types of replicons with respect to dimer resolution: small plasmids tend to use XerCD in a AS and PepA-depending manner; mid-size plasmids tend to use a self-encoded system, while the largest replicons tend to use XerCD in a FtsK-dependent manner, which becomes a rule above 250 kb.

### The R27 plasmid resolves dimers depending on FtsK

We analyzed how R27, a large plasmid (180.5 kb) isolated from pathogenic *Salmonella thyphi* (20), resolves dimers. Sequence analysis revealed a toxin-antitoxin system, which we inactivated and replaced by the *lacI* gene to ease stability studies (fig3A; Material and Methods). A GC-skew and a KOPS bias ran from replication origins to the opposite region, which contains an *xrs* devoid of predicted AS and a *matS* site. A gene encoding a site-specific recombinase of the Y-recombinase family (*Y-rec*) was also present. Stability assays shown that as the wild type R27, the *xrs* and the *y-rec* deleted variants of R27 were stably maintained in a *wt* strain (fig 3B). In contrast, the Δ(*y-rec*) variant was readily lost in strains inactivated for XerC, XerD or the ATPase activity of FtsK.

**Figure 3:**
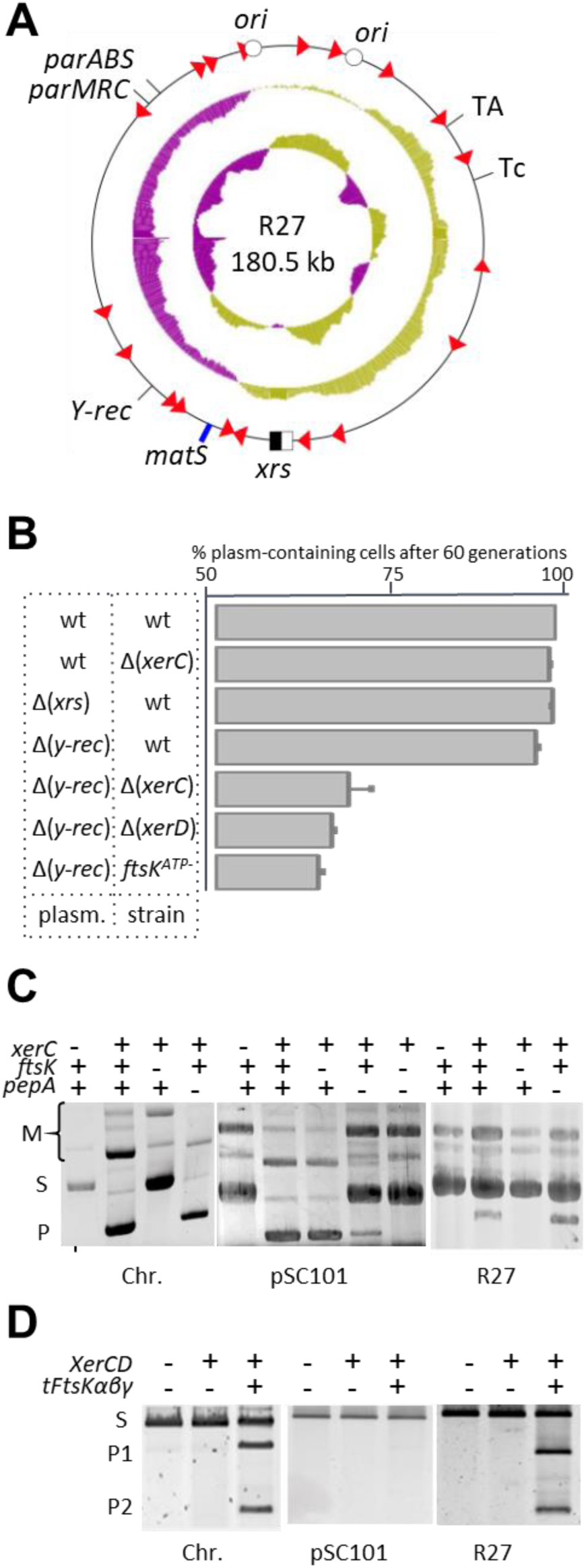
The R27 plasmid resolves dimers depending on FtsK. A) Map of R27 showing relevant DNA elements (see fig 1). The two replication origins (open dots) and the position of genes coding proteins relevant for plasmid maintenance (TA: toxin-antitoxin system; ParABS and ParMRC: partitioning systems; Tc: resistance to tetracycline; Y-rec: site-specific recombinase of the tyrosine family) are indicated. The middle circle shows a plot of the GC-skew and the inner circle a plot of the GC content. B) Loss of different R27 derivatives in relevant strains. Relevant genotypes of the plasmids (plasm.) and the strains are indicated. Strains were grown in serial cultures for 60 generation and plated on selective medium. C) Recombination between *xrs*. Plasmids carrying two *xrs* in direct repetition (see sup fig 3A) were introduced into the indicated strains and grown overnight before gel electrophoresis analysis. Positions of the starting plasmid (S), plasmid with deletion of the fragment separating the two *xrs* (P) and multimeric forms of these 2 plasmids (M) are indicated. The replicon from which the *xrs* originate are indicated below the gels (*xrs* from the chromosome (*dif*) and from plasmid pSC101 (*psi*) were used as controls). D) Activation of recombination by FtsK. The same plasmids as in C) were incubated with purified XerC and XerD and a variant of FtsK carrying a trimer of the C-terminal domain (tFtsKαβγ) as indicated before analysis by gel electrophoresis. Positions of the starting plasmid (S) and the two deletion products (P1 and P2) are indicated.

The R27 *xrs* is 28 bp-long and differs from known *xrs* in its XerC-binding site (fig S2A). It was less efficiently bound than the *dif* site by XerC in EMSA experiments (fig S2B), although complexes with XerD and with both XerC and XerD formed as efficiently as with *dif*. The R27 *xrs* was inserted with its flanking sequences as direct repetition in a multicopy plasmid to ease recombination studies (Material and Methods) and the resulting plasmid were used to transform chosen strains (fig 3C) and for *in vitro* recombination using XerC, XerD and an FtsK variant promoting hexamer formation and DNA translocation (tFtsKαβγ) (fig 3D). Comparing with equivalent plasmids containing the chromosome *xrs*, *dif*, or the pSC101 *xrs*, *psi*, a 28 bp-long *xrs* that mostly recombines depending on its AS (16, 29, 31), shown that the R27 *xrs* behaves as *dif* and was not affected by PepA inactivation but by FtsK inactivation. Taken together, our results show that R27 possesses two redundant systems to resolve dimers: a self-encoded Y-Rec system and an *xrs* recombining in an FtsK-dependent manner.

### Large replicons rely on FtsK-mediated control

As for the R27 *xrs*, we cloned *xrs* belonging to chosen replicons of different size (table 1) and assayed their capacity to recombine (Material and Methods ; Table 1). All cloned *xrs* recombined in an XerCD- dependent manner, showing they are active *xrs* (fig S3). Consistent with their association with ASs, plasmid pIGRW12 (5 kb) and R218 (114 kb) *xrs* recombined in a PepA-dependent manner. In contrast, *xrs* cloned from larger replicons, predicted not associated with AS, all recombined in an FtsK-dependent manner. FtsK-induced recombination assayed at a subset of the cloned *xrs* yielded the same conclusion (Table 1). These data validate our prediction of *xrs* and ASs and confirm that the *xrs* found in large secondary replicons recombine depending on FtsK.

**Table 1:** recombination at chosen *xrs*.

| replicon | size (kb) | Recombination |  |  |  |  |
| --- | --- | --- | --- | --- | --- | --- |
|  |  | wt | *xerC* | *ftsK* | *pepA* | *In vitro* |
| chr. | 4639 | + | - | - | + | + |
| pRAHAQ01 | 616 | + | - | - | + | ND |
| pEC-IMPQ | 324 | + | - | - | ND | + |
| pCFSAN | 294 | ND | ND | ND | ND | + |
| R478 | 275 | + | - | - | ND | + |
| pAPEC01R | 241 | ND | ND | ND | ND | + |
| pEQ1 | 239 | + | - | - | + | + |
| pHCM1 | 218 | + | - | - | ND | ND |
| R27 | 180 | + | - | - | + | + |
| RS218 | 114 | + | - | + | +/- | ND |
| pSC101 | 9 | + | - | + | +/- | - |
| ColE1 | 7 | + | - | + | - | - |
| pIGRW12 | 5 | + | - | + | - | - |

Recombination was monitored in vivo in a wild type (wt) or chosen mutant strains (for *xerC*, *xerD*, *pepA* or *ftsK*) transformed using reporter plasmids carrying a pair of *xrs*, cloned from the indicated replicon, in direct repetition, and in vitro using purified XerC, XerD and the tFtsK $\alpha\beta\gamma$ variant of FtsK, as in fig.2C and D, respectively (Material and Methods ). +: readily detected recombination products; -: no product detected; +/- : intermediate activity; ND: not done.

### KOPS biases and *matS* sites appear on large replicons

We next searched for KOPS and *matS* DNA sites, two DNA elements participating in the control of FtsK activity in the case of the chromosome (18). KOPS are recognized by FtsK and orient its activity, so that KOPS orientation pointing towards the *xrs* is important whereas KOPS occurrence is less or not important. Consistently, we did not find a significant enrichment of KOPS in replicons harboring *xrs* compared to replicons with no *xrs* (Material and Methods). To measure KOPS bias, we transformed the position of KOPS and their reverse motif as angles on a circle defined as one strand of the plasmid sequence centered on *xrs* (fig 4A; fig S4A). The mean angle of the KOPS positions thus points toward the position of the circle where KOPS are most concentrated (Material and Methods). Figure 4A shows that this mean direction of clustering of the KOPS and reverse KOPS motifs appear to point toward the middle of each halves of the replicon on both sides of the *xrs*. As plasmid size raises, sufficient KOPS were found to calculate significant clustering (41% of plasmids above 75 kb, yellow markers on fig 4A). Furthermore, when KOPS cluster in one half of the replicon, reverse KOPS tend to cluster in the opposite half, thus placing the *xrs* in the region of converging KOPS orientation (64 of the 78 plasmids with significant KOPS clustering; fig S4B). We conclude that KOPS biases tend to appear in replicons large enough to harbor multiple KOPS and that *xrs* tend to position in KOPS-converging regions.

**Figure 4:**
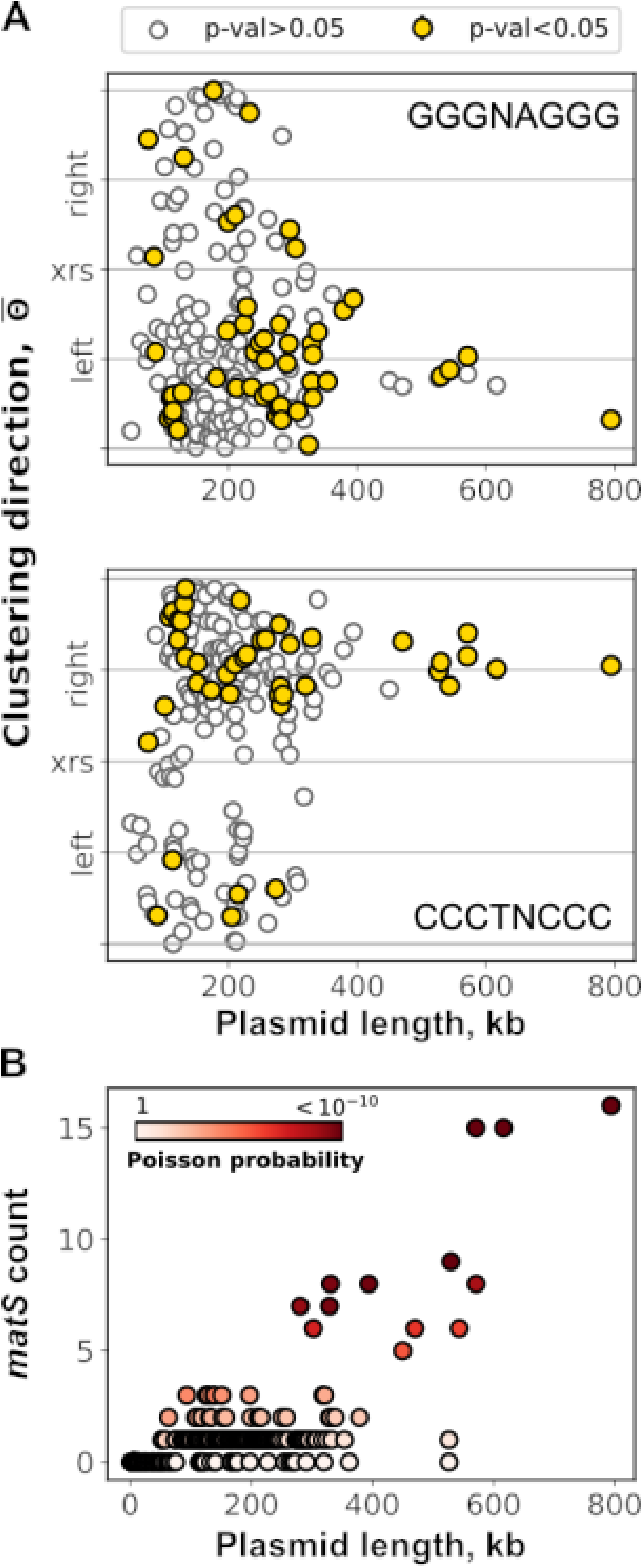
Large replicons tend to adopt a chromosome-like arrangement. A) KOPS biases appear in large replicons. Scatter plots of the mean angle (or direction) of the forward (GGGnAGGG, top) and reverse (CCCTnCCC, bottom) KOPS. Replicons for which the KOPS distribution significantly departs from uniformity (Material and Methods) are highlighted with gold colored circles outlined in black. Other replicons are symbolized by empty circles outlined in gray. Replicons with less than 3 KOPS were not considered in this analysis. B) *matS* are over-represented of in large replicons. Scatter plot of *matS* counts versus plasmid size. Each dot represents one replicon. The color code illustrates the probability of observing such counts in randomized sequences according to a Markov model with a chain length of 4 parameterized for each plasmid (Material and Methods).

The *matS* sites are palindromic 13 bp-long sequences degenerated at the three central positions (17). MatP is conserved across the bacteria included in this study, and the residues involved in the interaction with *matS* are identical (fig S4C). Detection of *matS* was performed using a position specific scoring matrix (Material and Methods ; fig S4D) constructed from the 23 *matS* sites identified on the *E. coli* chromosome (17). Most plasmids contained no *matS*, which was expected from the size of *matS*. Replicons larger than 100 kb tend to contain at least one *matS* and replicons larger than 300 kb contained at least two *matS* with few exceptions (fig 4B). To determine whether or not *matS* are enriched in these replicons, we measured *matS* occurrence in series of recoded plasmids of the same length and base composition (Material and Methods), allowing to calculate a Poisson probability of how likely it is to observe by chance the actual number of *matS* in the real plasmid sequence (fig 4B). We concluded that replicon above 300 kb are significantly enriched in *matS*. Importantly, replicons enriched in *matS* belong to different families, showing that *matS* appearance occurred independently in these different replicons.

## Discussion

A mean of dealing with the dimerization of circular replicons is certainly mandatory, as dimers and larger multimers pose a threat to the stable maintenance of replicons, interfering with their segregation and, in some cases, replication (12). For replicons replicating in theta mode, dimer resolution relies on site-specific recombination, which must be controlled so that dimers are better resolved than created. We have shown that the mode of control used primarily depends on replicon size. Replicons larger than 250 kb all contain a Xer recombination site (*xrs*) recombining in an FtsK-dependent manner as the chromosome-borne *xrs*, *dif*. This contrasts with replicons of other size range, which either use a different control of XerCD/*xrs* recombination in the case of small high copy-number plasmids or a self-encoded recombination system. We conclude that replicons above 250 kb have to adopt a chromosome-like control of dimer resolution. Consistently, the largest secondary replicons tend to acquire a KOPS-converging zone around their *xrs*, as well as *matS* sites, controlling FtsK and MatP activities, respectively, during segregation of the main chromosome.

The control of XerCD/*xrs* recombination in small plasmids involves a global structuration of the dimeric form into which the accessory sequences (AS) flanking the *xrs* get together by slithering over the complete length of the plasmid (fig S1) (16). Other recombination systems use the same kind of control (32). This mechanism obviously depends on plasmid size and may not be efficient enough in the case of large replicons, e.g. the efficiency of recombination by the RES/*res* system of transposon γδ drops two orders of magnitude when the two *res* sites are separated by 100 kb (33). The prevalence of FtsK-mediated control in large replicons may thus be primarily due to the poor efficiency of other kinds of control. The prevalence of 28 bp long *xrs* devoid of AS in large replicons also suggest that recombination systems other than XerCD/*xrs* do not easily acquire FtsK-mediated control. This may appear surprising as the Cre/*loxP* system from bacteriophage P1 has been shown to substitute functionally to the chromosome *dif* site in an FtsK-dependent manner (29). Consistently, the *loxP* site is located opposite to the replication origin in the plasmidic form of P1 and the P1 genome shows replichore organization from *ori* to *loxP*, with detectable GC skew and KOPS bias (34). However, only chromosome dimer resolution by Cre/*loxP*, not recombination as such, depends on FtsK, allowing the Cre/*loxP* system to play other roles. A strict dependence on FtsK may thus be selected in the case of large replicons and may be easily acquired only in the case of XerCD/*xrs*.

It is generally admitted that chromids originate from plasmids growing in size and acquiring features of the main chromosome, as bases and codon composition and core genes (5, 6). This occurred independently in bacterial orders, as judged by the different replication origins and initiator proteins. This evolution includes the adaptation of mechanisms involved in replicon maintenance. When known, chromids have the same copy number per cell as main chromosomes. Their replication is controlled so that termination is synchronous with that of the main chromosome (35–37), pointing to the post-replicative steps, unlinking and partitioning, as the most important to couple with the main chromosome and the cell cycle (12). We previously reported that the partitioning system used by replicons primarily depends on their size (28). In enterobacteria, secondary replicons above 200 kb all possess a ParAB*S* partition system, whereas smaller replicons may have any or no partition system. An equivalent step-size applies to the acquisition of FtsK-dependent dimer resolution, coupling the unlinking of sister replicon to cytokinesis in the case of main chromosomes. It thus appears that at least in enterobacteria, the 200 to 300 kb size range corresponds to a transition above which replicons cannot rely only on plasmid-like maintenance mechanisms and must acquire chromosome-like ones.

Consistently, few replicons are in this size-range and very few above (fig 2). We suspect that these replicons are selected for their very low copy number because, in addition to their metabolic load, their size can interfere with the normal dynamics of the main chromosome, as suggested by simulation studies (28). Their low copy number and large size then condition the maintenance required: efficient ParABS system for partitioning and integration of the unlinking step with cell division.

## Acknowledgments

We thank Laurence Van Melderen for help with TA annotation, Sophie Schbath for help with statistics and the members of the Genome Dynamics group of the CBI for helpful discussions.

## Funding

This work was supported by the Centre National de la Recherche Scientifique, the Université Toulouse III - Paul Sabatier and the Agence Nationale de la Recherche (Paris, FR), grant number ANR-14-CE10-0007 to FC and EPCR.

Supplementary data are available at NAR onlineS

---

**Figure S1.**
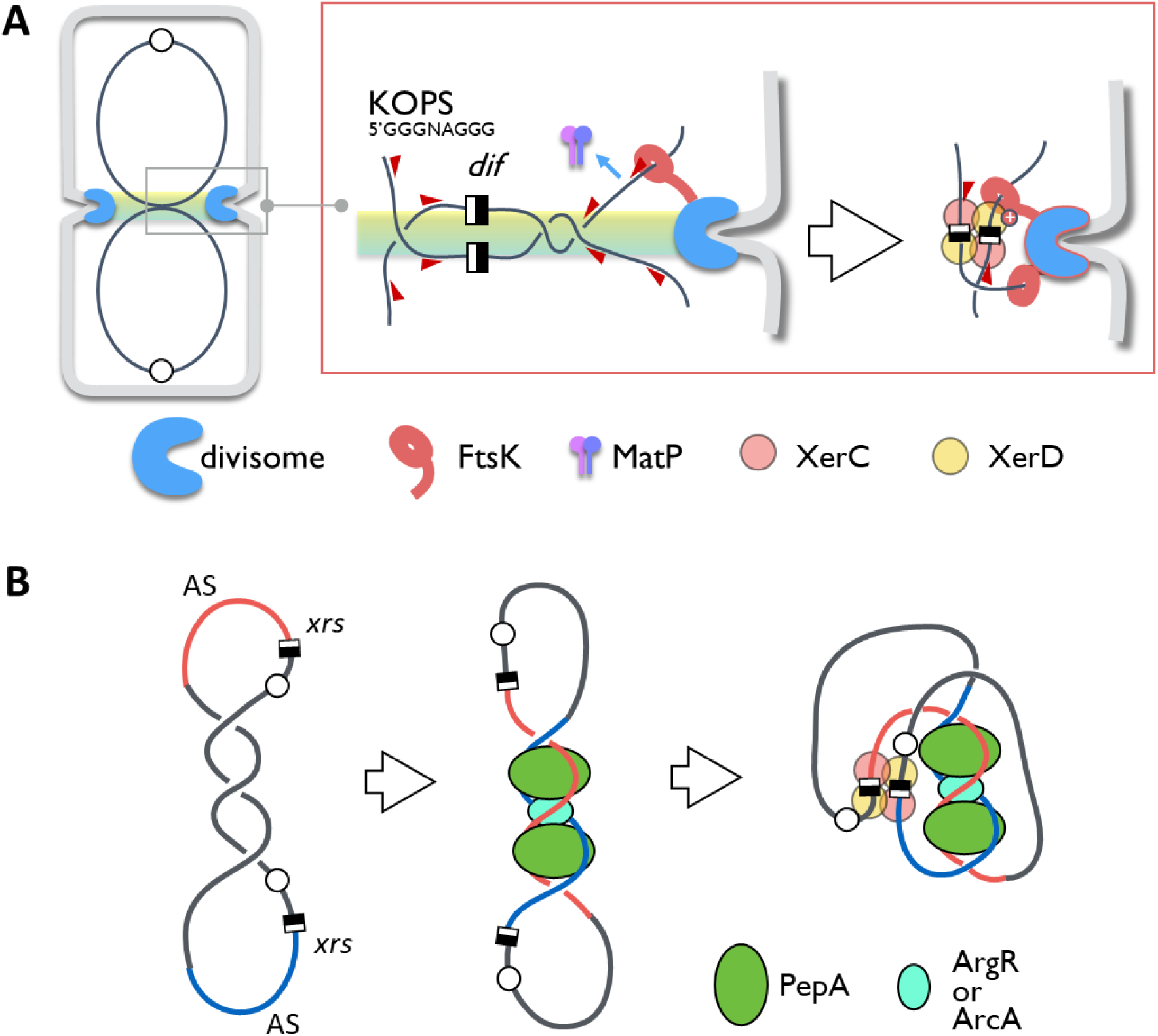
Control of Xer recombination. A) In the case of the chromosome, recombination occurs in pre-divisional cells at the onset of cell division (left drawing) and is controlled by the MatP and FtsK proteins. MatP dimers binds to *matS* sites scattered in the *ter* region (fig 1) and keep it in the mid-cell zone (yellow), allowing septum-borne FtsK hexamers to load onto *ter* DNA and translocate DNA towards the *dif* site following the orientation of the KOPS motifs (red arrowheads). Upon reaching the XerCD/*dif* complex, FtsK induces recombination, resolving the chromosome dimer to monomers. B) Recombination between *xrs* carried by model small plasmid (i.e., pSC101 and ColE1) depends on DNA Accessory Sequences (AS) flanking the XerC binding site (red and blue). In a plasmid dimer, the two ASs get together by sliding into the plectonems created by negative supercoiling and are recognized by the PepA and either ArcA (pSC101) or ArgR (ColE1) proteins. This creates a structure trapping three DNA crossing, required to induce recombination resolving the dimer. This mechanism ensures recombination is favored between the *xrs* carried by a dimer.

**Figure S2:**
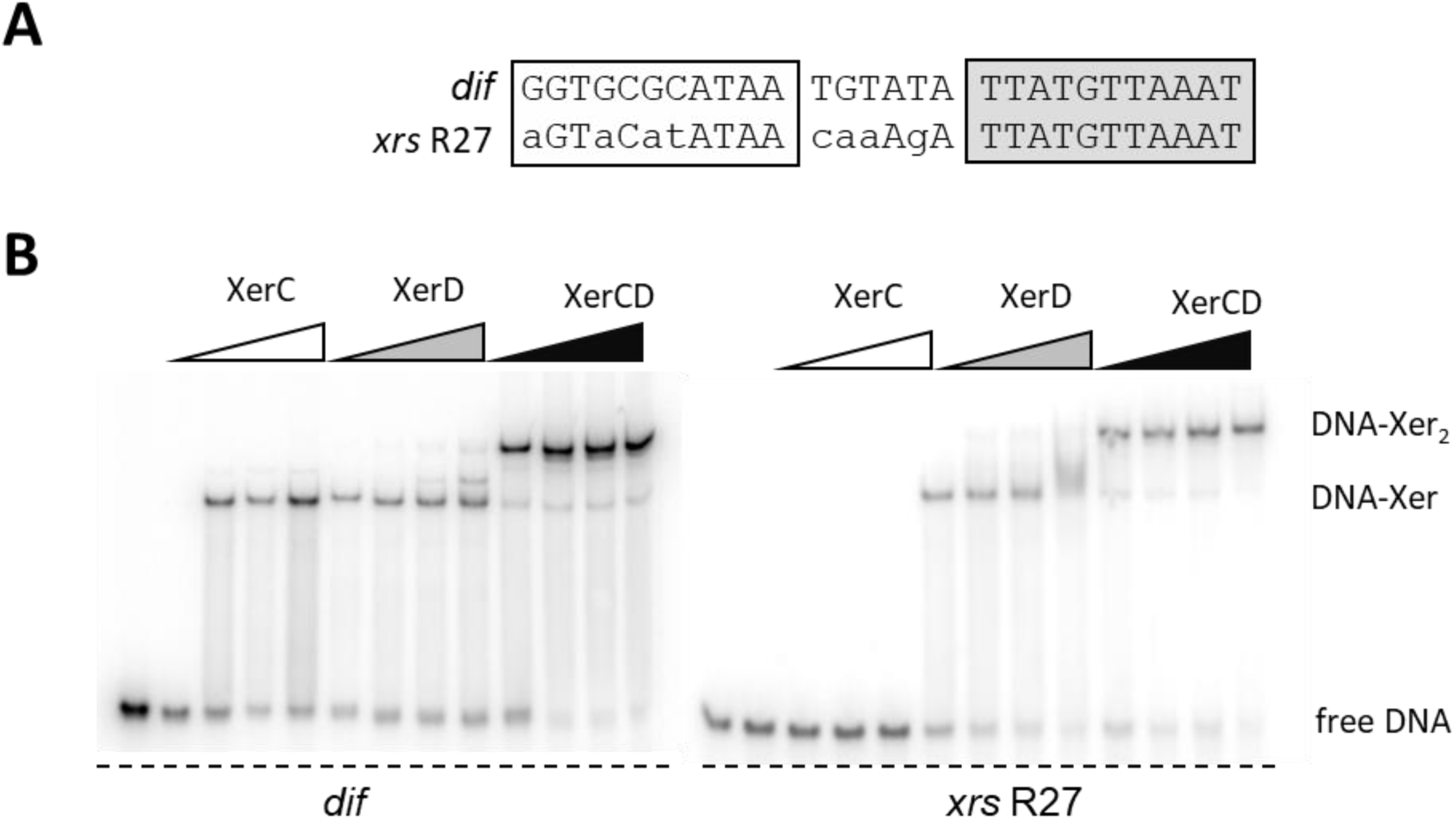
Interaction between XerCD and the *xrs* of plasmid R27. A) Sequence comparison between the *dif* site and the R27 *xrs*. XerC (left) and XerD (right) binding sites are boxed. Divergent bases are indicated in lower case. B) EMSA of *dif* or the R27 *xrs* with XerC and XerD. The radiolabelled DNA fragments are 28bp long. XerC and XerD ranged from 0.2 mM to 0.8 mM. Free DNA, DNA complexed to one recombinase (DNA-Xer), and DNA complexed to two recombinases (DNA-Xer2) are indicated.

**Figure S3.**
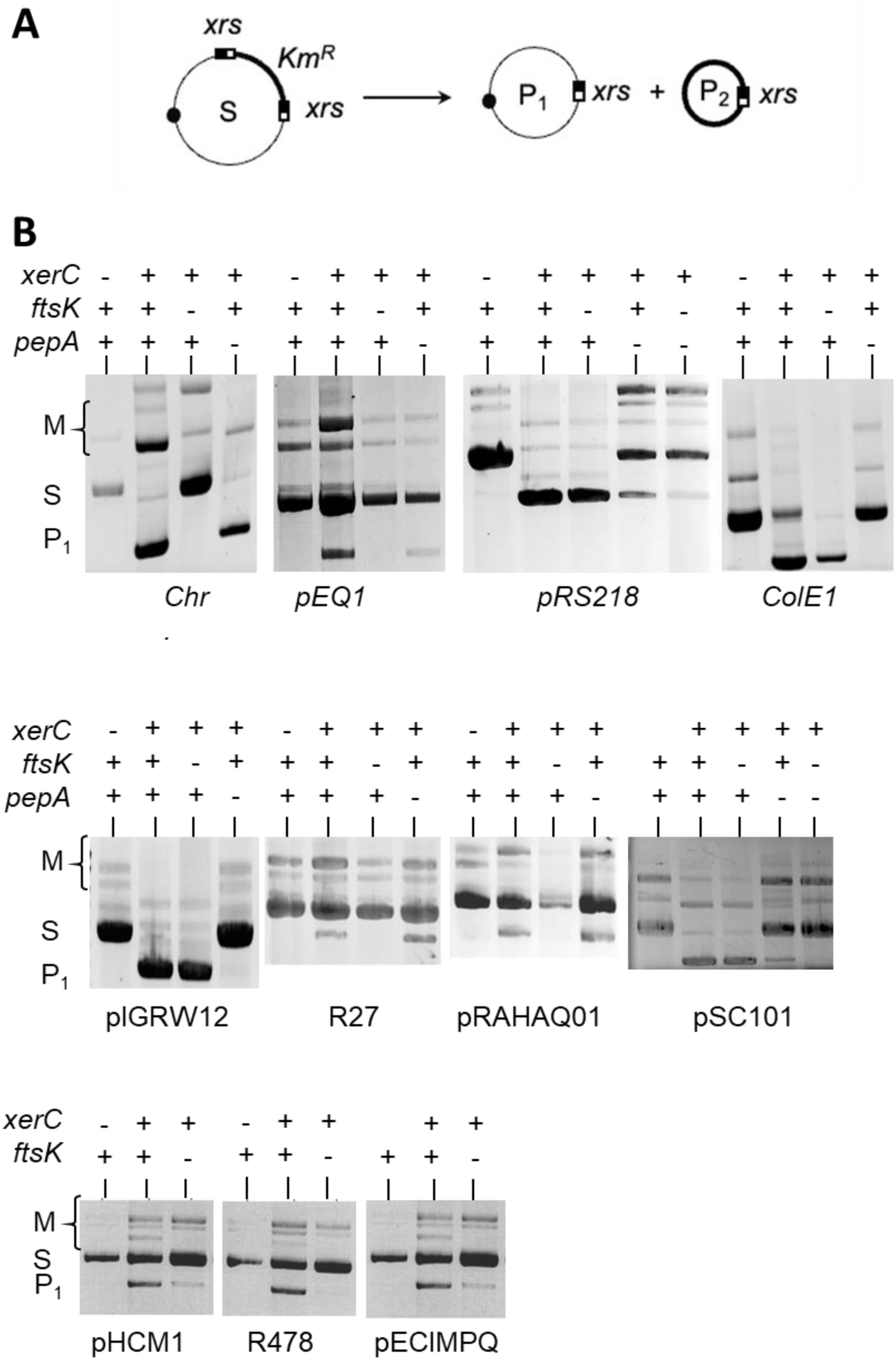
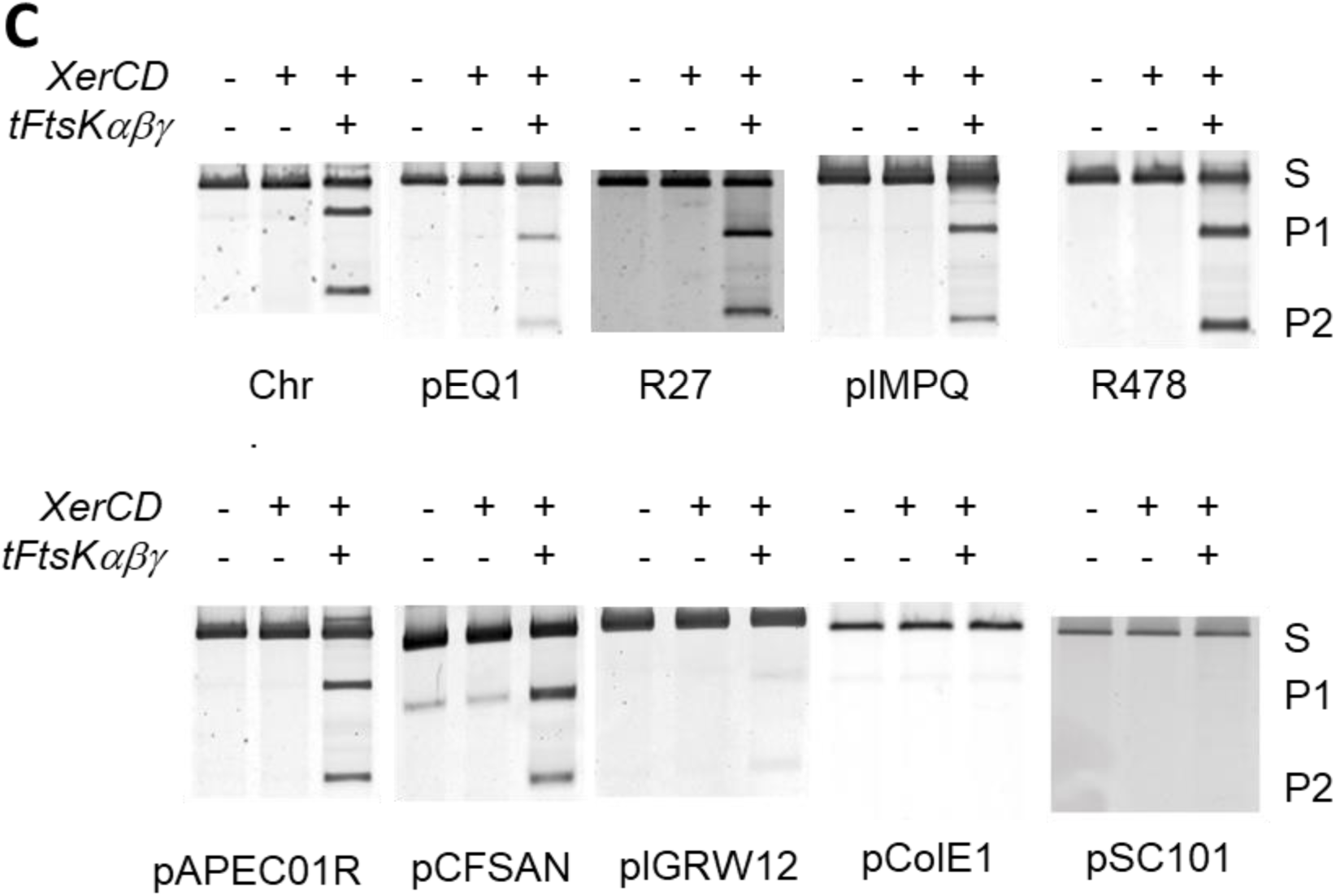
Recombination at cloned *xrs*. A) Recombination assay use plasmids carrying directly repeated *xrs*. Recombination deletes the intervening sequence yielding two recombination products (P1 and P2). P2 is not replicative, thus not retrieved in *in vivo* assays. B) *In vivo* recombination. Plasmids were transformed into the relevant strains carrying the indicated mutation (Material and Methods). Transformants were grown before plasmid extraction and analysis by gel electrophoresis. Position of the starting plasmid (S) and P1 product (P1), and their multimeric forms (M) are indicated. C) In vitro recombination. Plasmids were incubated with XerCD and tFtsKαβγ, a variant of FtsK containing a trimer of the C-terminal domain, as indicated, then analysed by gel electrophoresis. Positions of the starting plasmid (S) and the P1 and P2 recombination products are indicated

**Figure S4:**
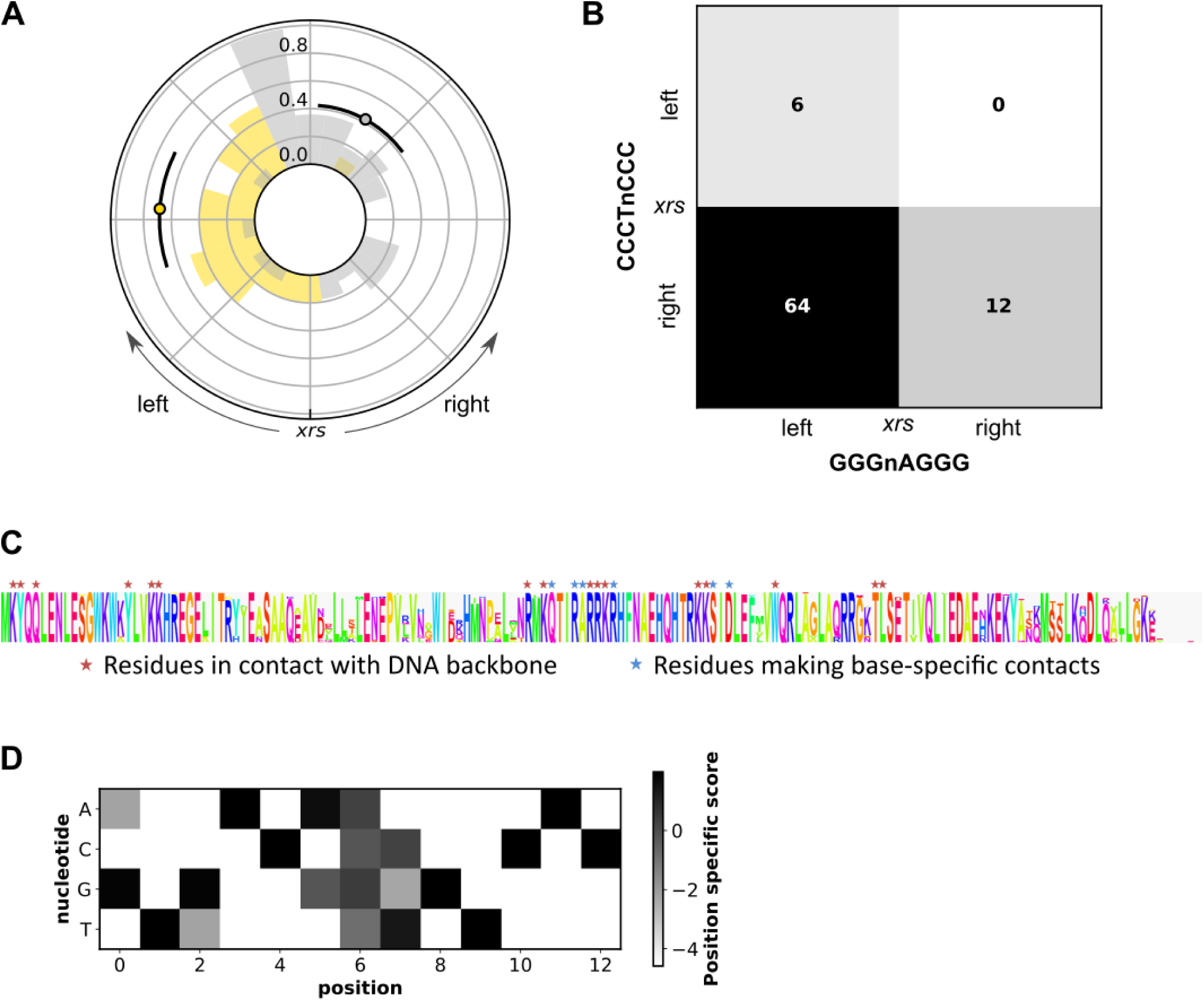
KOPS instances tend to cluster on opposite half of plasmids and *matS* sites are expected to be the same in all bacterial genomes of the collection. A) Circular histogram of KOPS in their forward (grey bars) and reverse (gold bars) orientation over the sequence of the largest plasmid of the collection (NCBI id: NC_014838). The circular sequence is orientated so that the xrs site is situated at the middle of the sequence, with a right and left half dominated by the presence of the KOPS motif or its reverse complement, respectively. B) Contingency table showing the number of plasmids with clustering of the KOPS motif in its forward and revers orientation in the left or right half of the DNA molecule. Only plasmids with significant clustering of the KOPS motif are considered in this table. C) Protein alignment of 402 MatP sequences annotated in the 427 unique genomes associated with the 914 plasmids of the collection. D) Position Specific Scoring Matrix derived from the 23 *matS* sites identified on the chromosome of *E. coli* K12 MG1655 (Mercier et al., 2008).

## References

1. Smillie, C., Garcillán-Barcia, M.P., Francia, M.V., Rocha, E.P.C. and de la Cruz, F. (2010) Mobility of plasmids. Microbiol. Mol. Biol. Rev., 74, 434–452.

2. Shintani, M., Sanchez, Z.K. and Kimbara, K. (2015) Genomics of microbial plasmids: classification and identification based on replication and transfer systems and host taxonomy. Front Microbiol, 6, 242.

3. Smalla, K., Jechalke, S. and Top, E.M. (2015) Plasmid detection, characterization and ecology. Microbiol Spectr, 3.

4. de Toro, M., de la Cruz, F. and Garcillán-Barcia, M.P. (2015) Plasmid Diversity and Adaptation Analyzed by Massive Sequencing of Escherichia coli Plasmids. In Tolmasky, M.E., Alonso, J.C. (eds), Plasmids: Biology and Impact in Biotechnology and Discovery. American Society of Microbiology, pp. 219–235.

5. diCenzo, G.C. and Finan, T.M. (2017) The Divided Bacterial Genome: Structure, Function, and Evolution. Microbiol Mol Biol Rev, 81, e00019–17.

6. Fournes, F., Val, M.-E., Skovgaard, O. and Mazel, D. (2018) Replicate Once Per Cell Cycle: Replication Control of Secondary Chromosomes. Front Microbiol, 9, 1833.

7. Hall, J.P.J., Botelho, J., Cazares, A. and Baltrus, D.A. (2022) What makes a megaplasmid? Philos Trans R Soc Lond B Biol Sci, 377, 20200472.

8. Garcillán-Barcia, M.P., Alvarado, A. and de la Cruz, F. (2011) Identification of bacterial plasmids based on mobility and plasmid population biology. FEMS Microbiol. Rev., 35, 936–956.

9. Dolejska, M. and Papagiannitsis, C.C. (2018) Plasmid-mediated resistance is going wild. Plasmid, 99, 99–111.

10. Alekshun, M.N. and Levy, S.B. (2007) Molecular Mechanisms of Antibacterial Multidrug Resistance. Cell, 128, 1037–1050.

11. Marchetti, M., Capela, D., Glew, M., Cruveiller, S., Chane-Woon-Ming, B., Gris, C., Timmers, T., Poinsot, V., Gilbert, L.B., Heeb, P., et al. (2010) Experimental evolution of a plant pathogen into a legume symbiont. PLoS Biol., 8, e1000280.

12. Cornet, F., Blanchais, C., Dusfour-Castan, R., Meunier, A., Quebre, V., Sekkouri Alaoui, H., Boudsoq, F., Campos, M., Crozat, E., Guynet, C., et al. (2023) DNA Segregation in Enterobacteria. EcoSal Plus.

13. Pesesky, M.W., Tilley, R. and Beck, D.A.C. (2019) Mosaic plasmids are abundant and unevenly distributed across prokaryotic taxa. Plasmid, 102, 10–18.

14. Harrison, P.W., Lower, R.P.J., Kim, N.K.D. and Young, J.P.W. (2010) Introducing the bacterial ‘chromid’: not a chromosome, not a plasmid. Trends in Microbiology, 18, 141–148.

15. Crozat, E., Fournes, F., Cornet, F., Hallet, B. and Rousseau, P. (2014) Resolution of Multimeric Forms of Circular Plasmids and Chromosomes. Microbiol Spectr, 2.

16. Colloms, S.D. (2013) The topology of plasmid-monomerizing Xer site-specific recombination. Biochem. Soc. Trans., 41, 589–594.

17. Mercier, R., Petit, M.-A., Schbath, S., Robin, S., El Karoui, M., Boccard, F. and Espéli, O. (2008) The MatP/matS Site-Specific System Organizes the Terminus Region of the E. coli Chromosome into a Macrodomain. Cell, 135, 475–485.

18. Stouf, M., Meile, J.-C. and Cornet, F. (2013) FtsK actively segregates sister chromosomes in Escherichia coli. Proc. Natl. Acad. Sci. U.S.A., 110, 11157–11162.

19. Reijns, M., Lu, Y., Leach, S. and Colloms, S.D. (2005) Mutagenesis of PepA suggests a new model for the Xer/cer synaptic complex. Mol. Microbiol., 57, 927–941.

20. Sherburne, C.K., Lawley, T.D., Gilmour, M.W., Blattner, F.R., Burland, V., Grotbeck, E., Rose, D.J. and Taylor, D.E. (2000) The complete DNA sequence and analysis of R27, a large IncHI plasmid from Salmonella typhi that is temperature sensitive for transfer. Nucl. Acids Res., 28, 2177–2186.

21. Colloms, S.D., Sykora, P., Szatmari, G. and Sherratt, D.J. (1990) Recombination at ColE1 cer requires the Escherichia coli xerC gene product, a member of the lambda integrase family of site-specific recombinases. J. Bacteriol., 172, 6973–6980.

22. Cornet, F., Louarn, J., Patte, J. and Louarn, J.M. (1996) Restriction of the activity of the recombination site dif to a small zone of the Escherichia coli chromosome. Genes Dev., 10, 1152–1161.

23. Diagne, C.T., Salhi, M., Crozat, E., Salomé, L., Cornet, F., Rousseau, P. and Tardin, C. (2014) TPM analyses reveal that FtsK contributes both to the assembly and the activation of the XerCD-dif recombination synapse. Nucleic Acids Res., 42, 1721–1732.

24. Crozat, E., Meglio, A., Allemand, J.-F., Chivers, C.E., Howarth, M., Vénien-Bryan, C., Grainge, I. and Sherratt, D.J. (2010) Separating speed and ability to displace roadblocks during DNA translocation by FtsK. EMBO J, 29, 1423–1433.

25. Grainge, I., Lesterlin, C. and Sherratt, D.J. (2011) Activation of XerCD-dif recombination by the FtsK DNA translocase. Nucleic Acids Res., 39, 5140–5148.

26. Collaboration, T.A., Price-Whelan, A.M., Lim, P.L., Earl, N., Starkman, N., Bradley, L., Shupe, D.L., Patil, A.A., Corrales, L., Brasseur, C.E., et al. (2022) The Astropy Project: Sustaining and Growing a Community-oriented Open-source Project and the Latest Major Release (v5.0) of the Core Package*. ApJ, 935, 167.

27. Schbath, S. and Hoebeke, M. (2011) R’MES: A Tool to Find Motifs with a Significantly Unexpected Frequency in Biological Sequences. In *Advances in Genomic Sequence Analysis and Pattern Discovery*, Science, Engineering, and Biology Informatics. WORLD SCIENTIFIC, Vol. Volume 7, pp. 25–64.

28. Planchenault, C., Pons, M.C., Schiavon, C., Siguier, P., Rech, J., Guynet, C., Dauverd–Girault, J., Cury, J., Rocha, E.P.C., Junier, I., et al. (2020) Intracellular Positioning Systems Limit the Entropic Eviction of Secondary Replicons Toward the Nucleoid Edges in Bacterial Cells. Journal of Molecular Biology, 432, 745–761.

29. Capiaux, H., Lesterlin, C., Pérals, K., Louarn, J.M. and Cornet, F. (2002) A dual role for the FtsK protein in Escherichia coli chromosome segregation. EMBO Rep, 3, 532–536.

30. Blakely, G. and Sherratt, D. (1996) Determinants of selectivity in Xer site-specific recombination. Genes Dev., 10, 762–773.

31. Colloms, S.D., Alén, C. and Sherratt, D.J. (1998) The ArcA/ArcB two-component regulatory system of Escherichia coli is essential for Xer site-specific recombination at psi. Mol. Microbiol., 28, 521– 530.

32. Grindley, N.D.F., Whiteson, K.L. and Rice, P.A. (2006) Mechanisms of Site-Specific Recombination. Annual Review of Biochemistry, 75, 567–605.

33. Higgins, N.P., Yang, X., Fu, Q. and Roth, J.R. (1996) Surveying a supercoil domain by using the gamma delta resolution system in Salmonella typhimurium. J Bacteriol, 178, 2825–2835.

34. Łobocka, M.B., Rose, D.J., Plunkett, G., Rusin, M., Samojedny, A., Lehnherr, H., Yarmolinsky, M.B. and Blattner, F.R. (2004) Genome of Bacteriophage P1. Journal of Bacteriology, 186, 7032–7068.

35. Rasmussen, T., Jensen, R.B. and Skovgaard, O. (2007) The two chromosomes of Vibrio cholerae are initiated at different time points in the cell cycle. EMBO J, 26, 3124–3131.

36. Du, W.-L., Dubarry, N., Passot, F.M., Kamgoué, A., Murray, H., Lane, D. and Pasta, F. (2016) Orderly Replication and Segregation of the Four Replicons of Burkholderia cenocepacia J2315. PLOS Genet, 12, e1006172.

37. Dubarry, N., Willis, C.R., Ball, G., Lesterlin, C. and Armitage, J.P. (2019) In Vivo Imaging of the Segregation of the 2 Chromosomes and the Cell Division Proteins of Rhodobacter sphaeroides Reveals an Unexpected Role for MipZ. mBio, 10, e02515–18.

