## Supplementary material for "The pathway to resolve dimeric forms distinguishes plasmids from megaplasmids in Enterobacteriaceae": sup file 2

### xrs previously characterized

Sequences are shown with their XerC-binding site left and their central region isolated (middle 6 to 8 bp). Names of their replicon is given. These *xrs* were used to start the search. Retreived xrs are shown in sup file 1.

| | | | |
|---|---|---|---|
| GGTGCGCATAA | TGTATA | TTATGTTAAAT | *E. coli* (*dif*) |
| GGTGCCGACAA | CGGATG | TTATGGTAAAT | ColA |
| GGTGCGTACAA | CGGGAG | TTATGGTAAAT | ColE3 |
| GGTGCACGCAA | CAGATG | TTATGGTAAAT | pJHCMW1 |
| GGTACCGATAA | GGGATG | TTATGGTAAAT | CloDF13 |
| GGTGCGCGCAA | GATCCA | TTATGTTAAAC | pSC101 |
| GGTGCGCGTAA | TGAGACG | TTATGGTAAAT | NTP16 |
| GGTGCGTACAA | TTAAGGGA | TTATGGTAAAT | ColE1 |

### ArgR binding site – containing accessory sequences (most divergent exemples)

Potential ArgR binding sites are bolded. The seven first ASs were previously characterized and used to start the search.

>**ColE1**, gi|9507253:3708-3938 Plasmid ColE1, complete sequence
ACCCTCTGGT**TGCAT**AGGT**ATTCA**TACGGTTAAAATTTATCAGGCGCGATCGCGCAGTTTTTAGGGTG
GTTTGTTGCCATTTTTACCTGTCTGCTGCCGTGATCGCGCTGAACGCGTTTTAGCGGTGCGTACAATT
AAGGGATTATGGTAAAT
>**ColA**, gi|9507286:1868-2096 Plasmid ColA, complete sequence
TTCACCTGGC**TGCAT**GGTT**ATGCA**GTCGGGTAAAATTTATCAGGTGCGTTCGGGGCGGTTTACCGGGT
GGTTTGTTGCCGGTTTTACCTGTCTTCCGCCCCGAACGCAGTGAACACGTCCGGAGCGGTGCCGACAA
CGGATGTTATGGTAAAT
>**ColE3**, gi|487267:160-371 Escherichia coli plasmid ColE3-CA38 DNA, replicon region encoding Rep protein
ATCGCTTTCG**TGAAT**AGTT**ATGCA**GGCCCCTGAAAACGATTCTGACGCGTTTTTTCGGTTTTGCCTGG
TGTTTTCCTGTCTTTTTGCGTTTTTTGCGTCAGAACGCGTCTGAGGGCGTTTTAAGGGGTGCGTACAA
CGGGAGTTATGGTAAAT
>**CloDF13**, gi|10955253:4773-5001 Escherichia coli K-12 P678-54 plasmid CloDF13, complete sequence
**TGCAG**TTGCG**TGCAT**AGGC**ATGCA**TTAGGATAAAATTTACCGGGCGCGTTTCCGGCGGTTTTCCGGGT
GGGTTGTTGCTTGTTTTATCCCGTAGCCGCCGGAAACGCCGCCAGTGCCTTACTGGCGGTACCGATAA
GGGATGTTATGGTAAAT
>**NTP16**, gi|9507429:2141-2348 Plasmid NTP16, complete sequence
TTCACTTTCA**TGCAT**AGCT**ATGCA**GTGAGCTGAAAGCGATCCTGACGCATTTTTCCGGTTTACCCCGG
GGAAAACATCTCTTTTTGCGGTGTCTGCGTCAGAATCGCGTTCAGCGCGTTTTGGCGGTGCGCGTAAT
GAGACGTTATGGTAAAT
>**pJHCMW1**, gi|19774233:185-485 Klebsiella pneumoniae plasmid pJHCMW1, complete sequence
TGCACTTGCG**CGCAT**ACTC**ATGCA**TGCCGTAAAAACAGAGCCTGCGCGTTTCTGGCGGGTTTTCGGGT
GGTTTGTTGCCTGTTTTACCGGTTTCCCGTCAGAAACGCCCTGAGGGCCTCTCAGGCGGTGCACGCAA
CAGATGTTATGGTAAAT
>**pIGRW12**, gb|EF088686.1|:2104-2404 Escherichia coli plasmid pIGRW12, complete sequence
TGCATCTGGC**TGCAT**GACT**ATGCA**CCAGTGTAATATTCAGCGGCTGCGTTTCTGGTGTTTTTTCGTTA
TGTCTGTCGCCACTTTCTCGCGTCTGGCACCGGAAACGCAGCCAACCCCGTTTCAGCGGTGCGGGCAA
CGGATGTTATGGTAAAT

>**pAm08CD7339xrs1**, gb|GQ149345.1|:4884-6784 Escherichia coli plasmid pAm08CD7339, complete sequence
TCCATTGCGG**TGCAT**GGCT**ATTCA**TGGCCTCACAATTCATCAGAGGCGTTTTGCGGGATTTTAGGCGG
TAAAGAGGTGCGGTTCTGTGGCTTAGCCGCCCAAAACGCGCGTAGAGCGTCTCTGGCGGTGCGGGAAA
TGAGGCGTTATGGTAAAT

>**pAm08CD7339xrs2**, GQ149345.1:c6784-6457 Escherichia coli plasmid pAm08CD7339, complete sequence
GGTATGTCCG**TGCAT**AGCC**ATGCA**GTCGCGCCAGAATCATCAGGGGCGTTTTTCTCGATATGGGCAC
AGAAAGGTGTGTGTTTCAGGCCGGAGGCCTCAGAAACGCCCTCAGAGGCTCTGAGGCGGTGCACGCAA
CGGATGTTATGGTAAAT

>**pUMNturkey5_5** gb|JRQA01000059.1|:666-966 Escherichia coli strain UMNturkey5 plasmid pUMNturkey5_5, whole genome shotgun sequence
TTCATTTTCG**TGCAT**GGCT**ATGCA**TACACGCAAAACTTATCAGCGCGATTCCGGGCGGTTTTCGGTGA
GAAGACGGGCTCTTTTGTGGCTCAGGACGGAAAAACGCCCTGAGCGCGTCTGAGCGGTGCGCCCAAGG
GATGTTATGGTAAAT

>**PCN061p1**, gb|CP006637.1|:c1434-1134 Escherichia coli PCN061 plasmid PCN061p1, complete sequence
TTCACTTCTG**TGCAT**AGCC**ATGCA**GGCGCGCAGAAATCATCAGGCGCGTTTTTTCACGATATGGACAG
GGAAAGGTGCCGGAAACGCAGAGGACGCGGCAGAAACGCGCTGAGGGCCTCTCAGGGGGTGCGGGCAA
GGGATGTTATGGTAAAT

>**pNPO1**, gb|KF992024.1|:c1692-1392 Escherichia coli strain W1058 plasmid pNPO1, complete sequence
GGCATTTCCG**TGCAT**AGCC**ATGCA**GACGCGCAGAAATCATCAGGAACGTTTTTCGGGGTTTTCCCCGG
GGAAAGGTGCCGGAAATCGGCATGAGGCCGCAGAAACGCGCTGAGAGCGTCTTAGGCGGTACCGATAA
GGGATGTTATGGTAAAT

>**pAPEC-O78-3**, gb|CP010318.1|:1-76 Escherichia coli strain O78-789 plasmid pAPEC-O78-3, complete sequence
TTCAGTTTCG**CGCAT**AACT**ATGCA**TGAGGTTAAAATTTACCAGGCGCGATCGCGGCAGTTTTTCGGGT
GGTTTGTTGCTGTTTTTACCTGTCTGCTGCCGTGATCGCGCTGAACGCGTTTAAGCGGTACGCGCAAT
GCGACGTTATGGTAAAT

>**pCGB40**, gb|JQ776504.1|:3818-4118 Escherichia coli strain CGB40 plasmid pCGB40, complete sequence
GTCAGTCAGG**TGCAT**GGTT**ATGCA**TGGGGCTGAAAATCACGGTATGCGATTCTGAGCGGTTTTCGGTG
AGAAGACGGGGGTTTTTATGGCTCAGGACGTAAAACCGCCCTGAGCGCGTTTCAGCGGTGCGCGTAAT
GACGCGTTATGGTAAAT

>**pMNCRE44_5**, gb|CP010881.1|:c97299-96999 Escherichia coli strain MNCRE44 plasmid pMNCRE44_5, complete sequence
TTCGGTTGAG**TGCAT**ATCC**ATTCA**TAGGGTAGATTCTTAAGTCGCGTTTCTGGTGTTCATTTTCGGGT
GGTTTGTTACTTGTTTTACCGGGGATATGCCAGAAACGCGCTGAGTCAGTCTGGGCGGTGCGCGTAAT
GCGGCGTTATGGTAAAT

>**pSE11-5**, AP009245.1:1610-1765 Escherichia coli SE11 plasmid pSE11-5 DNA, complete sequence
ACCATCTGGT**TGCAT**AGGT**ATTCAT**GCGGTTAAAATTTATCAGGCGCGATCGCGGTAGCTTTTCGGGT
GATTTGTTGTTGGTTTTGGCTGACTGCCGCCCCGTTCGCGGCGAAGCTGTCCGGGGCGGTGCGGGCAA
CAGATGTTATGGTAAAT

>**p1303_5**, CP009169.1:c4099-3946 Escherichia coli 1303 plasmid p1303_5, complete sequence
TTCAGTTTCG**TGCAT**AGTC**ATGCA**ACGCCCTGAAAACGATCCTGACGCATTTTTCCGGTTTTCCTGGG
GGTAAACATTTCTTTTTGCTGTGCCTGCGTCAGAATCGCGCTCAACGCGTTTTAATGGTGCGTACAAT
TAAGGGATTATGGTAAAT

>**pLST424C-10**, NC_019092.1:688-841 Escherichia coli plasmid pLST424C-10, complete sequence
TGCACCTGGC**TGCAT**AGCC**ATGCA**TCCGGGTATGATTTATCCGGTGCGTTTCTGGCGGGTTTTCGGGT

GGTTTGTTGCCGCTTTTACCGGTTATCCGTCAGAAACGCGCTGAGTCAGTCTGGGCGGTGCGCGTAAT
GGGACGTTATGGTAAAT

>**pEC278**, AY589571.1:2858-3011 Escherichia coli strain 278B plasmid pEC278, complete sequence

TTCACCCGAC**TGCAT**AGCC**ATGCA**GCCGGGTGAAATTTATCTGGCGCAGAATGCGCGGTTTAGCGGAG
AGTTTGCTGCCGGTTTTACCGGGCTGACGCCGCCATTTGTTCCGAACCCGTCCGGGGCGGTGCTGATA
AGGGATGTTATGGTAAAT

>NC_013954.1:2075-2402 Erwinia pyrifoliae strain Ep1/96 complete plasmid pEp2.6

GGCATTCTGC**TGCAT**AACC**ATGCA**TACAGGCAAAAGCGCCTGAGGGCGTTTCTGACGAATTTTCTCCA
GGAAGACAAGGTTTTTTACGGCTTACGGCTTCAAAGGGCTCTGAGTGCGTCTCAGGCGGTGCGGGCAA
CGGGTGTTATGGTAAAT

>NC_017444.1:1713-1913 Erwinia sp. Ejp617 plasmid pJE05, complete sequence

GTTAGTCCGG**CGCAT**AACG**ATTCA**ACCGCTCAGAACTTACCAGAGCGATTCTGAGCGGTTTTCAGTGG
GGGATATGCGGGAGTTTTGGCGGTTTGACCTCAAAATCGTGCTGAGGGGCTCTCAGGCGGTTCGCGCA
ACGGGTGTTATGGTAAAT

>NZ_CP012166.1:1243-1443 Enterobacter hormaechei subsp. oharae strain 34978 plasmid p34978-2.725kb, complete sequence

GTCATTCTGG**TGCAT**AGTC**ATGCA**TGCCGTTAAAATTTAACAGGAGCGTTTCTGGCGGGTTCCGGGGT
GGTTTGTTGTGGTTTTTGGTCATGGTTCCGTCAGAAAAGCGCTGAGTGCGTCTGAGGCGGTGCGGGCA
ACGGATGTTATGGTAAAT

>NZ_CP016445.1:c2903-2703 Edwardsiella piscicida strain S11-285 plasmid unnamed1, complete sequence

TGCAGTGTCG**TGAAT**GGCT**ATGCA**ACGCCGCCAGAATCATCCTGCGCGATTTTGGCCGTCTTGGGGGT
ACGAAGCTGCGTCTTTTGCCGATAACGCCTCAGAATCGCGCTCAGCGCGTCTGAGGTGGTGCGCCGAG
CGGACGTTATGGTAAAT

>NC_020212.1:c370-170 Serratia marcescens WW4 plasmid pSmWW4, complete sequence

CACGCCGGGT**TGAAT**AACC**ATTCA**CGCTGCCAAACGGCCTGTATGCTATTTTGCGGGCGTTTTAGCGC
AGAAGTGAGGAGGTTTTGTGGCTTCCGCCTCAGAAGCGCGCTGAGGGTGCCTGAGGCGGTGCGGGAAA
CGGATGTTATGGTAAAT

>NZ_CP008845.1:1133-1333 Klebsiella michiganensis strain M1 plasmid pKOXM1D, complete sequence

TGCAGTGATG**TGAAT**AAGTG**ATGCA**TATGCGTAAAATAAATCAGGTGCGTTTCTGGTGGTTTTCGGTG
TACTTGCCACCACCTTTTGCCCGTCAGCCATCAGAAACGCCCTGAGTGCCTCCGGCGCGGTGCGGGTA
AGAAGACGTTATGGTAAAT

>NZ_CP011316.1:c3777-3669 Klebsiella pneumoniae subsp. pneumoniae strain 234-12 plasmid pKpn23412-4, complete sequence

CGCACTTGCG**TGCAT**ACTC**ATGCA**TGACGTGAAAACAGAGCTAGCGTGTTTTTGGCTGATTTTTTGAT
AGTTTGTTGCCTGTTTTACCAGTTTCCCGTCAGAAACACCCTGAGGCCGTTTGGGCGGTGCGTACAAT
TAGGGTGTTATGGTAAAT

>NZ_CP011434.1:1-78 Salmonella enterica subsp. enterica strain YU39 plasmid pYU39_4.2, complete sequence

TTCACTCCTG**TGCAT**AGCC**ATGCA**GACGCGCAGAAATCATCAGCGCGTTTCTGAGCGGTTTTCAGGGG
GTAAGATGTGTTCTTTTGCAGGTGAGGTCGTAGAAACGCCCTGAGAGCCTCTCAGGCGGTGCACGCAA
CAGATGTTATGGTAAAT

>NC_009652.1:1295-1495 Klebsiella pneumoniae subsp. pneumoniae MGH 78578 plasmid pKPN6, complete sequence
GTCAGTCTGG**TGCAT**AACG**ATTCA**TGCCCGTAAAACGCCCTGGAGCGATTTTGAGGCGTTTTCAGGGG
GTAAGACGTAGGATTTAATGGGTTACGCCCCAGAATCGTTCTGAGGCCGTTTTAGCGGTGCGTGTAAT
GACGCGTTATGGTAAAT
>NZ_FO818639.1:7427-7627 Xenorhabdus bovienii str. CS03 plasmid XBC_p, complete genome
CACGGGGGAA**TGCAT**AATT**ATGCA**TTCGTTGAAAAACGGCTGTGCGCGATTTTGCGGGTTTTTGGAGG
GAAAGAGCGTGTGTTTTACCGGGTTGAGGCGAAAACTGCGCTGAATGGCGTACAGGCGGTTCTGGCAA
CGGCTGTTATGTTAAAT
>NZ_CP011627.1:12470-12623 Klebsiella oxytoca strain CAV1374 plasmid pCAV1374-14, complete sequence
TGCGGTGGTG**TGCAT**AGCC**ATGCA**TATGCGTAAAATAAATCAGGTGCGTTTCTGGCGGGTTTTCGGGT
GGTTTGTTGCCTGTTTTACCGGTTTCCTGCCAGAAACGCCCTGAGGCCGTTTTCGCGGTGCGCGTAAA
AAGGCTTTATGTTAAAT
>NZ_CP009366.1:c38191-37991 Yersinia enterocolitica strain WA plasmid, complete sequence
GCAGCCAGGC**TGCAT**AGGT**ATTCA**TGCGGTTAAAATTTATCGGGTGCGATCGCGATAGTTTTTCGGGG
AGTTTGTTGCTCTTTCTGCCTGTTTACTGCTGTGATCGCGCTAAACGCATTTCTGCGGTGCGTTAAAT
CCATCTTATGTTAAAT
>NZ_CP017187.1:c48843-48690 Enterobacter cloacae complex 'Hoffmann cluster III' strain DSM 14563 plasmid pDSMZ14563, complete sequence
AGCACTTTTG**TGCGT**TTGT**ATGCA**TTGAGGTAAAATTCATCTCCTTCGTTTCTGGCGGGTTTGTTGGT
GGGTTGTTGCCTGTTTCACCCGGTTCCTGTCAGAAACGCGCTCCGGCCGTCTGAGCTGTGCGCGTAAT
GAAGCATTATTGTAAAT
>NZ_CP014127.1:c57954-57802 Pantoea agglomerans strain FDAARGOS_160 plasmid unnamed2, complete sequence
GAGCTCTCAC**TGCAT**AGCC**ATGCA**GATGCGCCGAAATCATCAGGAGTATTTTAAGACGTTTAAGCAGA
GAAAGGTGCCGGAAACGGCCCTCAGGCCGCAGAAACGCGCTGAGCGCGCCTGCGGGGGTGAAGGCAAG
CACTGTTATGTTAAAG

### ArcA binding site – containing accessory sequences

Potential ArgR binding sites are bolded. The first AS was previously characterized and used to start the search.

>**pSC101**, gi|10955533:6582-6810 Salmonella enterica subsp. enterica serovar Typhimurium plasmid pSC101, complete sequence
CAAACCTGAAGCCGGATCTGCGATTCTG**ATAACAAACT**AGCAACACCAGAACAGCCCGTTTGCGGGCA
GCAAAACCCGTACTTTTGGACGTTCCGGCGGTTTTTTGTGGCGAGTGGTGTTCGGGCGGTGCGCGCAA
GATCCATTATGTTAAAC
>**pECO-b75**, CP009861.1:23-178 Escherichia coli strain ECONIH1 plasmid pECO-b75, complete sequence
ATGAGGCGAAACTGAAAACGAGATTTCA**TTAACAAATA**AGCAACATAAAAAACGGTGGAAAACGCCCA
GGAAAACATGATCTTTTGAGCGGATTTTTTAGATCCGACAATGAACGGTGATCCGGTGGTGCCGATAA
CGTCCATTATGTTAAAT
>NC_010695.1:1400-1556 Erwinia tasmaniensis strain ET1/99 complete

plasmid pET09
ACCGGCTGAAACCGGATCCGCGATTCTG**ATAACAAGCC**GGCAACACCAAAACAGCCCGTTTTTGGCCG
GCAACACCCGCGCTTTTGGACGTTCCGGCGGGTTTTGGCGATGAGTGGTGTTGCAGTGGTGCGCGCAA
GAACTCTTATGTTAAAT
>NZ_LN890526.1:29396-29596 Salmonella enterica subsp. enterica serovar Weltevreden genome assembly 99_3134, plasmid : 3
TTGACCTGAAATTAGAAACGACTTTTTA**GTAACAAATT**AGTAACATTAAAAGCTATGCTAAATTACTA
GGTAAAGTTGATCTTTTGAGGTGTTTACTCCGATATTGGTGCCAAAGTGGATCGAGTGGTGCCGATAA
CGACCATTATGTTAAAT
